# A Tumor Suppressor-Regulated Cell Cycle Derived Gene Signature is Prognostic of Recurrence Risk in Prostate Cancer

**DOI:** 10.1101/397331

**Authors:** Constantin Georgescu, Joshua M. Corbin, Sandra Thibivilliers, Zachary D. Webb, Yan D. Zhao, Jan Koster, Kar-Ming Fung, Adam S. Asch, Jonathan D. Wren, Maria J. Ruiz-Echevarría

## Abstract

**Background:** The clinical behavior of prostate cancer (PCa) is variable, and while the majority of cases remain indolent, 10% of patients progress to deadly forms of the disease. Current clinical predictors used at the time of diagnosis have limitations to accurately establish progression risk. Here we describe the development of a tumor suppressor regulated, cell-cycle gene expression based prognostic signature for PCa, and validate its independent contribution to risk stratification in several radical prostatectomy (RP) patient cohorts.

**Methods:** We used RNA interference experiments in PCa cell lines to identify a gene expression based gene signature associated with *Tmeff2,* an androgen regulated, tumor suppressor gene whose expression shows remarkable heterogeneity in PCa. Gene expression was confirmed by qRT-PCR. Correlation of the signature with disease outcome (time to recurrence) was retrospectively evaluated in four geographically different cohorts of patients that underwent RP (834 samples), using multivariate logistical regression analysis. Multivariate analysis were adjusted for standard clinicopathological variables. Performance of the signature was compared to previously described gene expression based signatures using the SIgCheck software.

**Results:** Low levels of Tmeff2 mRNA significantly (p<0.0001) correlated with reduced disease-free survival (DFS) in patients from the Memorial Sloan Kettering Cancer Center (MSKCC) dataset. We identified a panel of 11 TMEFF2 regulated cell cycle related genes (TMCC11), with strong prognostic value. TMCC11 expression was significantly associated with time to recurrence after prostatectomy in four geographically different patient cohorts (2.9≤HR≥4.1; p≤0.002), served as an independent indicator of poor prognosis in the four RP cohorts (1.96≤HR≥4.28; p≤0.032) and improved the prognostic value of standard clinicopathological markers. The prognostic ability of TMCC11 panel exceeded previously published oncogenic gene signatures (p=0.00017).

**Conclusions:** This study provides evidence that the TMCC11 gene signature is a robust independent prognostic marker for PCa, reveals the value of using highly heterogeneously expressed genes, like *Tmeff2*, as guides to discover prognostic indicators, and suggests the possibility that low *Tmeff2* expression marks a distinct subclass of PCa.

## INTRODUCTION

Cancer of the prostate (PCa) is the second leading cause of cancer death in male Americans. The clinical behavior of PCa is variable, and while the majority of PCa cases remain indolent, 10% of patients progress with aggressive metastatic disease and subsequent emergence of therapy-resistant PCa (1, 2). In current practice, clinical variables including Gleason score, tumor stage, and PSA levels are used at the time of diagnosis to predict disease outcome (3, 4). However, these prognostic factors have limitations, resulting in significant rates of overtreatment, with associated comorbidities (5-7), and undertreatment, leading to disease progression and increased risk of PCa-specific mortality (8-10).

The clinical heterogeneity of PCa reflects, in part, a remarkable genomic heterogeneity (11-18). This suggests that disease stratification based on molecular features may be of prognostic value beyond standard clinicopathological variables, and aid in the clinical management of the disease, as is the case for other cancers, i.e. breast (19-21). Currently, several tissue-based molecular tests offer prognostic information for patients with PCa either before or after treatment. These are based on general features of malignancy, such as the Prolaris test (initially described by Cuzick et al.(22)), which incorporates information from 31 cell-cycle related genes, or on molecular features more specific for PCa (Decipher, Oncotype DX, ProMark, and ConfirmMDx tests (23-27)). In addition, recent work has outlined the existence of several molecular subtypes of PCa (28-31). Notably, in one of these studies, the molecular subtypes were defined by specific driver mutations or gene fusions that are essentially mutually exclusive and that were able to categorize up to 74% of the analyzed tumors (32). If shown to correlate with clinical behavior, these molecular subtypes could prove critical for the management and treatment of the disease. However, currently their prognostic value is not fully established, and a significant fraction of primary prostate cancers in the study, could not be categorized within these molecular subsets suggesting the existence of additional relevant molecular alterations.

High levels of variability in gene expression between tumors can be useful in identifying prostate and other cancers risk genes (33). We hypothesized that molecular subtypes of primary prostate cancers may exist that have gene expression patterns associated with changes in the expression of these highly variable genes. A recent report lists *TMEFF2* as one of the top 100 mRNA transcripts with the highest levels of inter-tumor variability in primary PCa tissues (34). TMEFF2 is an androgen regulated transmembrane protein mainly restricted to brain and prostate. Our studies in PCa demonstrate a role of TMEFF2 as a tumor suppressor (35-38). Furthermore, studies using limited numbers of clinical samples, reveal changes in the expression of *Tmeff2* with disease stage in PCa (39, 40) and gliomas (41), supporting an important role of *Tmeff2* in these diseases.

We have investigated the expression pattern of TMEFF2 in human prostate tissues and explored the potential of a TMEFF2 associated gene signature as a biomarker for disease prognosis. We report that low *TMEFF2* mRNA expression is associated with decreased disease free survival (DFS) in the MSKCC PCa dataset. Using transcriptional profiling of cell lines and publically available PCa clinical data, we have identified a low *TMEFF2* driven gene signature associated with poor clinical outcome, comprised of cell cycle related genes with increased gene expression. This study not only provides new insights into the clinical relevance of *Tmeff2* in cancer, but also specifies a group of cell cycle related proteins as prognostic and potential therapeutic targets.

## METHODS

### *TMEFF2* Expression data

*TMEFF2* mRNA expression in benign and malignant samples of PCa was interrogated using Oncomine Compendium of Expression Array data (42) in the following cohorts: Varambally et al. ((n=19; GSE3325; (43)), Vanaja et al. ((n=40; (44)), Grasso et al. ((n=122; GSE35988; (45)), and Taylor et al. ((or MSKCC; n=185; GSE21032; (46)).

### Validation cohorts

Four prostate cancer cohorts were used in this study to establish the prognostic value of the TMCC11 signature: MSKCC (46) (GSE21032); Cambridge (34) (GSE70768) and Stockholm (34) (GSE70769), are microarray datasets, and the TCGA PRAD (http://cancergenome.nih.gov/), an RNA sequencing cohort. Cancer samples for all cohorts were from RP specimens. Biochemical recurrence (MSCKK, Cambridge and Stockholm) or recurrence/progression (TCGA-PRAD) was the follow-up endpoint. Clinical, histopathological data and summary of the cohorts are listed in Tables 1 and S1 (Additional File 1).

**Table 1.**
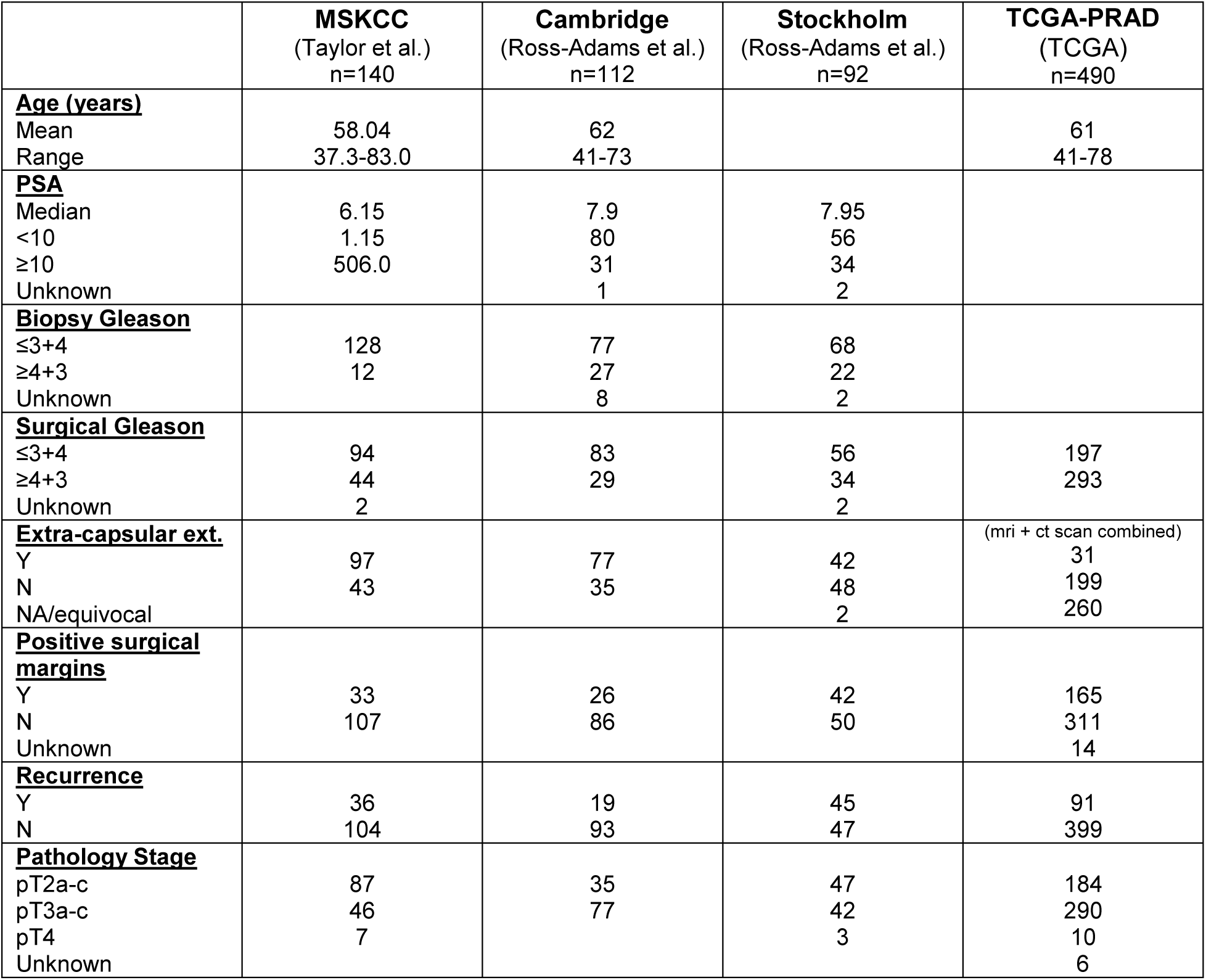
Clinical and pathological characteristics of the prostate cancer datasets used in this study. The table defines the characteristics of the samples used in this study for each of the datasets.

### Mammalian cell culture and treatment

The LNCaP and 22Rv1 cell lines were purchased from American Type Culture Collection (ATCC; Manassas, VA) and cultured as recommended. DHT (Sigma, Burlington, MA) was used at a concentration of 10nM. For TMEFF2 knockdown, LNCaP and 22Rv1 cells were transduced with pLKO.1 lentiviral vectors with antisense TMEFF2 sequences shTMEFF2-0 (TRCN0000073518), shTMEFF2-1 (TRCN0000073519) and shTMEFF2-2 (TRCN0000073521). See table S2 in Additional File 1 for sequences.

### RNA extraction and RNA-Seq

LNCaP cell expressing sh_TMEFF2 or the sh_scramble control were grown for 24 hrs in hormone-depleted media and stimulated with 10 nM DHT (or ethanol as vehicle control) for 24 hrs prior to harvesting for RNA extraction. Three biological replicates per sample were used. Total RNA was extracted with RNeasy mini kit (Qiagen, Waltham, MA) and cDNA was synthesized with SuperScript III First-Strand synthesis system (Life Technologies Inc, Carlsbad, CA). RNA integrity and quantity was assessed using the Agilent Bioanalyzer (Agilent Technologies, Santa Clara, CA). Raw 75bp paired-end sequences were generated from an Illumina NextSeq 500 sequencer (Illumina, San Diego, CA). Sequenced reads first underwent quality control with the FASTQC tool and then aligned to a contaminant genome to filter out reads which align to human ribosomal RNA, poly-A, poly-C, phiX virus or mitochondrial DNA sequence. The filtered reads were trimmed using Trimmomatic (47), as well as read clipping based on quality over a sliding window, retaining reads with a minimum length of 15 bp. Trimmed, filtered reads were pseudoaligned to the GRCh38 human reference transcriptome using kallisto version 0.42.3 (48), with enabled bias correction and 50 bootstrapping rounds. Expression values for 173259 unique transcripts were measured and transcripts with an average of 5 count per million (CPM) or less across all samples were removed from further analysis. To perform differential expression analysis (LNCaP-sh_TMEFF2 vs. LNCaP-sh_scramble control), CPM values were summarized at the gene level and normalized with the *R* packages (49) and DESeq2 (50) to identify significantly differentially expressed genes (DEGs) with fold change ≥ 1.5 and FDR-adjusted p-value ≤ 0.05. Data deposited in NCBI GEO under accession number GSE117180.

### Real-time polymerase chain reaction (RT-PCR)

Total RNA was extracted with RNeasy mini kit and cDNA was synthesized with iScript™ Reverse Transcription Supermix for RT-qPCR (BioRad, Hercules, CA). Quantitative RT-PCR was performed using the SsoAdvanced™ Universal SYBR^®^ Greenand gene specific primers (see Table S2, Additional File 1) on the Biorad CFX96™ Touch Real-Time PCR Detection System (BioRad, Hercules, CA). All RT-PCR experiments were performed under MIQE guidelines, using three biological replicates and two technical replicates.

### Western blotting

Cell lysates were prepared in RIPA buffer containing a protease inhibitor mixture and analyzed by Western blot as described before (38), using the following antibodies: TMEFF2 (HPA015587, Sigma) at a 1:1000 dilution; AR (sc-7305, Santa Cruz Biotechnology Inc, Dallas, TX) at a 1:1000 dilution; and Calnexin (ab22595; Abcam, San Francisco, CA) at a dilution of 1:4000.

### TMCC11 signature selection process

From the 25 genes initially identified as significantly upregulated (Log2 fold change ≥1.8, ≤3.1; FDR < 0.05) by DHT in the LNCaP-TMEFF2 knockdown cells, we selected the 21 top-ranking upregulated genes (Log2 fold change ≥2.0) (refer to figure S2 in additional file 1). We interrogated this 21 gene subset in the MSKCC dataset (n = 150) in cBioPortal (51, 52) and selected those genes (n=11; TMCC11) whose expression was upregulated in patients with low TMEFF2 mRNA expression (at least 4 of those patients) and that maintain a strong functional association as demonstrated using STRING (53) and IPA pathway analyses (additional file 1; Figure S2 and S3). Two other signatures were used for SigCheck analysis. The TMCC13 is a modified TMCC11 signature including two additional genes, E2F7 and GSG2 (from the TMEFF2 21 top-ranking upregulated genes; additional file 1; Figure S3), selected based on their individual prognostic values and lack of overlap with genes from the Cuzick (22) signature. TMCC3 consist of the CDC45, NCAPG and CLSPN genes and was selected from TMCC11 as the optimal subset in predicting time to BCR in the Stockholm dataset. For this purpose, the dependence of time to BCR on the signature gene expression was modeled using GLM cox regression, and the search for the best subset relied on elastic net regularization, a standard features selection procedure implemented in the R package glmnet.

### TMCC11 signature score development

Patients were divided in two categories (high and low) based on the TMCC11 gene signature, by calculating the mean expression over all the genes in the signature for each sample. The distribution for the population was calculated, and samples were included in the high group when their mean fell within the upper tertile (above the 67^th^ percentile) and in the low group when below the 67th percentile.

### Databases and statistics

Databases/platform used during this study: cBioportal (51, 52), Oncomine (42), the R2 genomic analysis and visualization platform (http://r2.amc.nl); the STRING database (53); and SurvExpress (54). The parameters used are referenced in the corresponding figure legends if applicable. For publicly available microarray or RNA-Seq expression data sets, the normalized expression data was downloaded from the Oncomine, cBioportal or R2 databases.

Hierarchical clustering of the TMCC11 signature genes (Euclidean distance with average linkage on zscore transformed expression values) on samples from the MSKCC dataset was performed in R2.

Data analysis was performed by non-parametric Wilcoxon multiple comparison test or Student t-test as indicated in figure legends. Statistical significance was defined as *P*<0.05 unless otherwise stated. Time-to-event outcomes were evaluated using Kaplan-Meyer analysis and survival-time differences compared using the log-rank test. Uni-, multi-variate and C-statistics were used to assess the independent effect of biomarker status on clinical outcome. Univariate hazard ratios and p-values were obtained using the Cox proportional hazard model. Multivariate analysis was performed using the Cox proportional hazard model. A stepwise model selection procedure coupled with Cox proportional hazard model was used to define the final model. The Harrell’s method was used to compute the concordance statistics. Covariates included in the multivariate models were Biopsy and/or surgical Gleason score, PSA, pathological T-stage, positive surgical margins and/or extracapsular extension, and were adjusted as follow: Gleason – High (≥4+3): Low (≤3+4); PSA – High (≥10):Low(<10); Path Stage –High(≥T3):Low(≤T2); Positive surgical margins −Y:N; Extracapsular extension (ECE) – Y:N. These analyses were conducted using SAS 9.4 and a p-value of less than 0.05 or 0.01 if indicated, was deemed statistically significant.

### Gene signature analysis with SigCheck

We analyzed the prognostic potential and specificity of the TMCC11 signature using the Bioconductor package SigCheck (55). This software allows comparison of a gene signature prognostic performance against random and known gene signatures. In a first analysis, we compared the prognostic power of the TMCC11 gene signature and a total of 253 oncogenic signatures available from literature. The prognostic power of a gene signature was quantified by the log-rank test p-value for the difference between the time to BCR in high versus low risk groups according to overall signature gene expression. Mean expression over all the genes in the signature for each sample was computed and high versus low expression was considered as over or below the 67th percentile respectively. Log-rank P-values for each signature were computed using the Stockholm13 (GSE70769), Cambridge13 (GSE70768) and MSKCC14 (GSE21034) datasets downloaded from the GEO website. In a second analysis, we comparatively assessed the superiority of the TMCC11 and the other 253 oncogenic signatures against randomly constructed predictors. For each signature under study, 10,000 signatures of the same number genes were selected at random and for each log-rank p-value scores of their predictive power were computed as described above. A bootstrap p-value was then determined as the proportion of random gene signatures scoring better than the original gene signature. Stockholm, Cambridge and MSKCC datasets were also used for this analysis. The code for the analysis is available upon request.

## RESULTS

### Low expression of TMEFF2 is associated with advanced disease and is prognostic of clinical outcome

The previously described cell growth inhibitory function of TMEFF2 in PCa (35-37) led us to determine the relationship of *Tmeff2* expression alterations to the clinicopathologic features of PCa. We first analyzed tumor associated changes in TMEFF2 expression by immunohistochemistry in PCa tissues (Figure S1A, additional file 1). TMEFF2 expression was higher in patients with localized disease as compared to non-tumor samples (not shown). However, when patients were stratified by tumor stage, TMEFF2 expression was significantly decreased in more advanced pathological stages (Figure S1B, additional file 1).

We then used Oncomine (42) to examine alterations of *TMEFF2* mRNA expression in publically available samples from PCa patients. Expression of *TMEFF2* mRNA is significantly increased in the primary tumors of patients with PCa when compared to normal tissue, in multiple independent datasets (Figure 1A). However, in samples from metastases and castration resistant prostate cancer (CRPC), the levels of *TMEFF2* mRNA are either unchanged or decreased compared to normal prostate, and significantly decreased (P<0.05) when compared to primary tumors (Figure 1A). These data suggest a negative correlation between *TMEFF2* mRNA expression and progression to the advanced stages of PCa.

**Figure 1.**
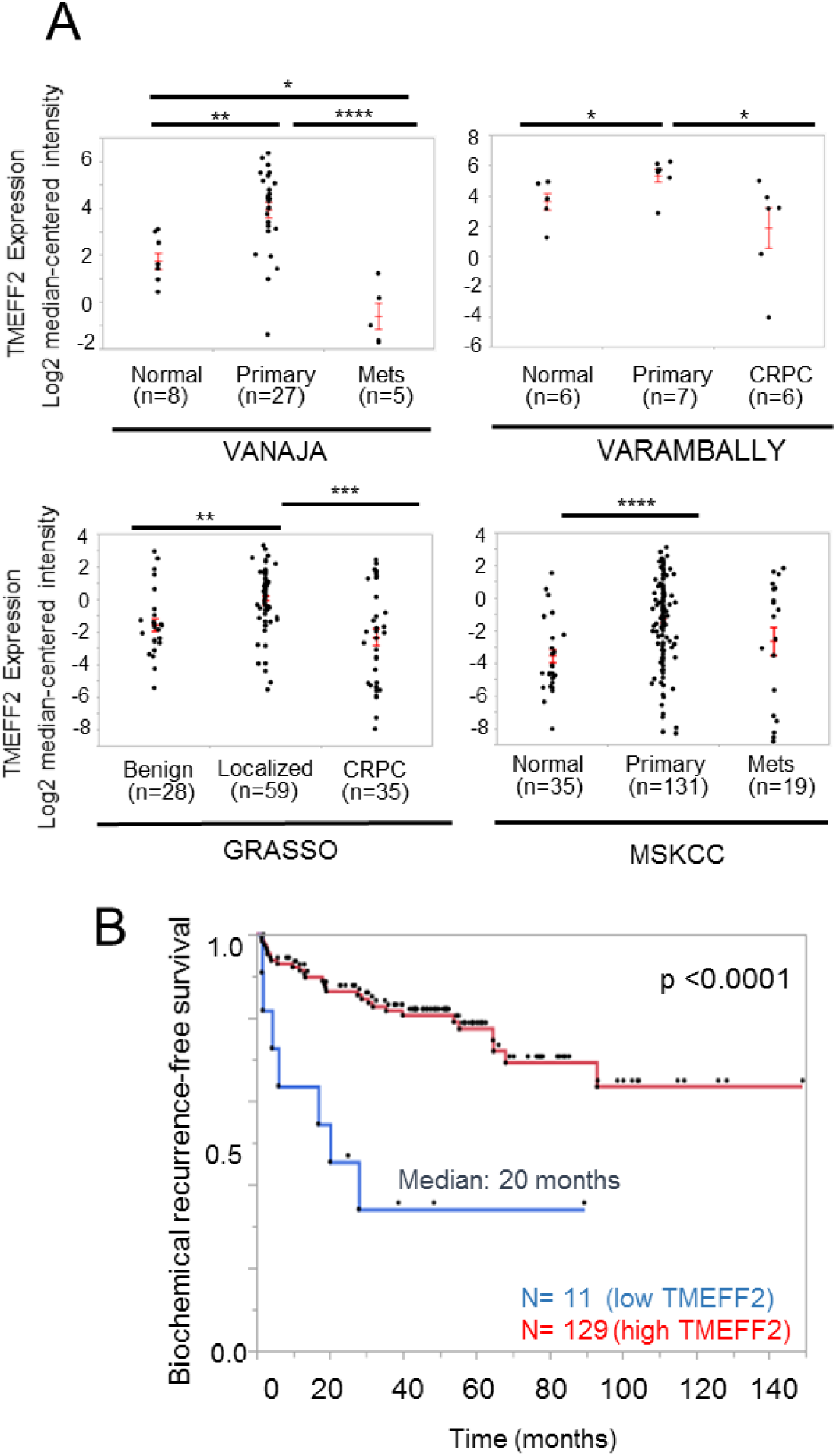
Low expression of TMEFF2 is associated with advanced disease and clinical outcome. 1A) Scatter plot showing TMEFF2 expression levels in normal, primary and metastatic/CRPC tissue from patients from different prostate cancer cohorts. Expression levels were obtained from Oncomine and are compared using a Wilcoxon multiple comparison test. **1B)** Kaplan-Meir plot showing survival of patients from the MSKCC prostate cohort with low TMEFF2 expression (n=11; lowest expression in the cohort) vs. the rest of the patients. *P<0.05; **P<0.01; ***P<0.001; ****P<0.0001.

Based on these observations, we analyzed the prognostic value of *TMEFF2* mRNA expression in the MSKCC dataset ((46); Table 1), a publically available human PCa dataset with clinical outcome data. Kaplan-Meier analysis demonstrated a significant (p<0.0001) correlation between *TMEFF2* levels and disease progression (assessed by biochemical recurrence, BCR). Patients with the lowest *TMEFF2* mRNA expression had higher BCR (20 vs. 110 months; Figure 1B). These findings underscore the clinical significance of *Tmeff2* in cancer.

### *TMEFF2* silencing in the LNCaP cell line increases androgen-driven expression of a group of cell-cycle related genes

*TMEFF2* is one of the top 100 mRNA transcripts with the highest levels of inter-tumor variability in patient samples from several publically available datasets ((34) and Table S1 Additional File 3). Such heterogeneity and the fact that low *TMEFF2* mRNA expression correlates with advanced disease, suggest that it may define a molecular signature with prognostic value. To begin understanding the molecular consequences of decreased *TMEFF2* expression and its potential to define a prognostic gene signature, we conducted *TMEFF2*-targeted RNA interference experiments. We found that *TMEFF2* silencing alters expression of androgen receptor (AR) targets, suggesting that previously reported TMEFF2 effects on growth (37) may be driven, in part, by TMEFF2-modulated AR-mediated expression of genes involved in cell cycle related processes. Using shRNA, we silenced expression of *TMEFF2* in LNCaP cells (Figure 2A, and additional file 1, figure S2A and S2B), a PCa cell line that expresses high levels of *TMEFF2* mRNA and protein. Using RNA-Seq, we identified a group of 25 nuclear genes that were moderately but significantly upregulated, in the context of *TMEFF2* silencing (Log2 fold change ≥1.8, ≤3.1; FDR < 0.05), by dihydrotestosterone (DHT) in the LNCaP-Tmeff2 knockdown cells as compared to control cells (transduced with scramble shRNA; Additional file1, Figure S2C). STRING pathway analysis (53) suggests that most of these genes are functionally associated (Additional file 1, Figure S2D) and belong to the DNA replication and cell cycle gene ontology categories.

**Figure 2.**
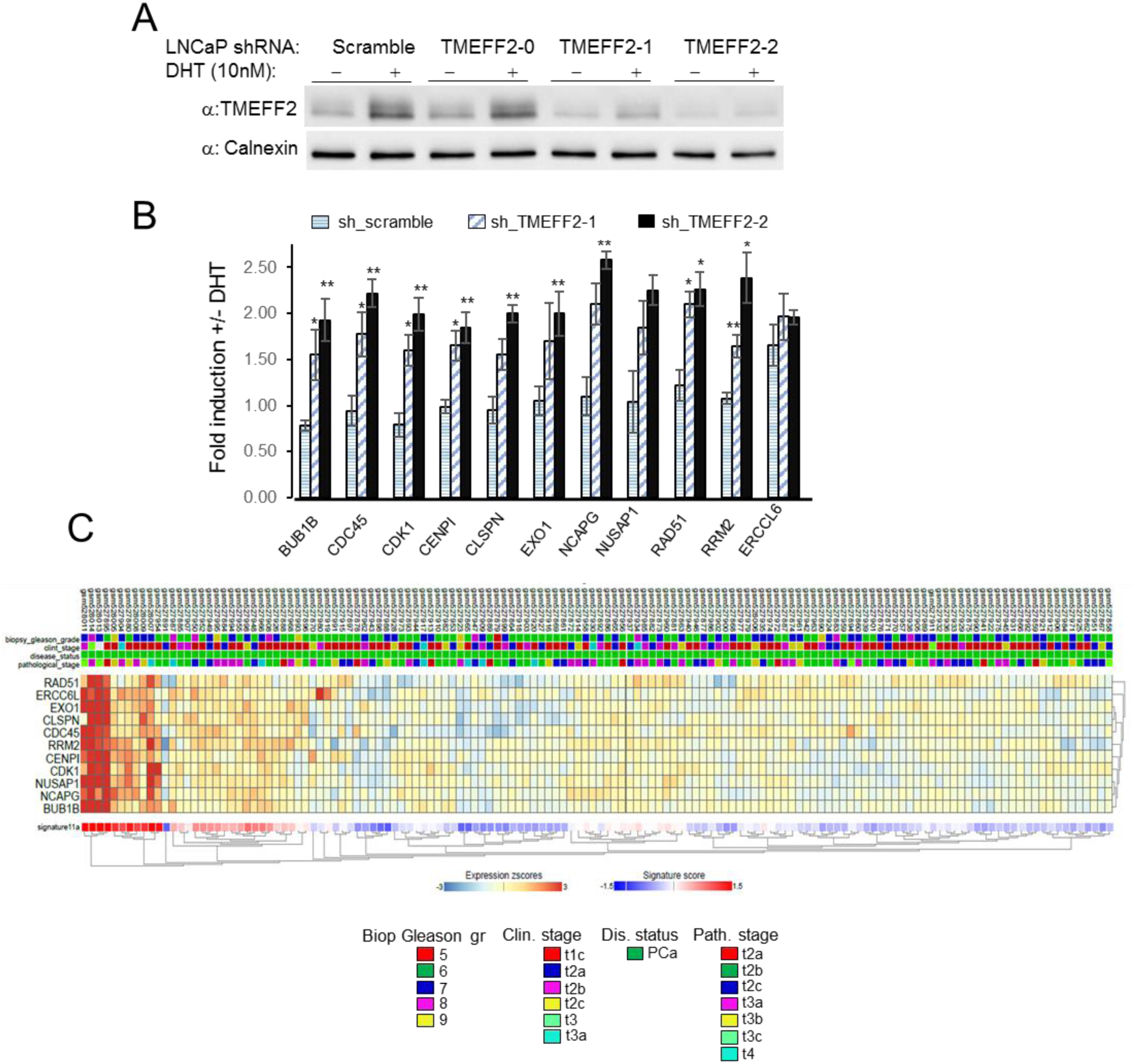
TMEFF2 silencing in PCa cells induces androgen-driven expression of cell cycle genes. 2A) Western Blot analysis to determine knockdown of TMEFF2 in LNCaP cells using three different Tmeff2 targeted sh-RNAs. Only sh_TMEFF2-1 and sh_TMEFF2-2 appreciably silenced Tmeff2 expression. Note that *Tmeff2* is an androgen regulated gene. Representative blot from >3 repeats. **2B)** qRT-PCR data in the LNCaP-sh_TMEFF2 cells confirming increased expression in response to androgen stimulation of the cell cycle genes selected for the TMCC11 signature. Data is the average of 3 independent repeats and was analyzed using T-test. Error bars correspond to s.e.m. **2C)** Clustering analysis of TMCC11 signature genes in the MSKCC cohort. Each column corresponds to individual patient. The status of some clinicopathological variables for each sample has been included in the figure at the top of the heatmap. *P<0.05; **P<0.01.

Out of the initial group of genes, we selected 11 (Additional file 1, Figure S3 and methods) referred to as the “*TMEFF2* modulated cell cycle 11 (TMCC11)” gene signature. qRT-PCR analysis in LNCaP cells confirmed that DHT mediated induction of the TMCC11 genes was significantly increased in LNCaP cells in which *TMEFF2* expression was low compared to control cells (Figure 2B). High expression of these genes with low TMEFF2 expression was also seen in patients samples from the MSKCC dataset (Additional file 1, Figure S3C). Clustering analysis of the TMCC11 signature genes in the MSKCC dataset indicate that expression of these genes is highly correlated (Figure 2C). These 11 genes are all tightly related to cell-cycle and DNA replication and repair processes (Additional file 1, Figure S3B). Moreover, silencing of *TMEFF2* in PCa cells affects cell cycle progression (Additional file 1, Figure S4) supporting the role of TMEFF2 in modulating expression of cell-cycle related genes.

In clinical samples from the Grasso (45) and MSKCC (46) datasets, the expression of the individual genes from the TMCC11 signature is significantly increased in CRPC and metastatic disease samples when compared to normal tissue, and inversely correlated with the expression of *TMEFF2* in the same samples (Additional file 1, Figure S5A and S5B). In addition, mRNA coexpression analysis using the PCa MSKCC and PRAD TCGA datasets indicates that these genes are significantly co-expressed (Additional file 1, Figure S6).

### The TMEFF2-modulated gene signature is an independent marker of recurrence after prostatectomy in multiple clinical datasets

Based on our results suggesting that loss of TMEFF2 often predates aggressive/metastatic disease, we postulated that the TMEFF2-modulated TMCC11 gene signature could have prognostic value. We evaluated this hypothesis using BCR as the clinical endpoint in the PCa MSKCC dataset (46) (Table 1, and Additional file 1, Table S1 and Figure S7, provide information on the samples). The MSKCC dataset includes a number of prostatectomy samples from patients with wide range of times to BCR as measured by increased levels of PSA. Individually, increased expression of each of the genes comprising TMCC11 was statistically significant (P< 0.01) in predicting BCR (Additional File 1, Table S4; for CLSPN p =0.0137). In Kaplan-Meier analyses, high expression of the TMCC11 signature was associated with a median time to progression of 55.39 months vs. greater than 150 months for patients with low expression of TMCC11 (log-rank P value = 1.11e-05; Figure 3A). These results indicate that the TMCC11 signature is a powerful predictor of aggressive PCa, segregating the tumors into high and low-risk groups based on time to BCR. We obtained similar results using the SurvExpress (54) database for analysis (Additional File 1, Figure S8).

**Figure 3.**
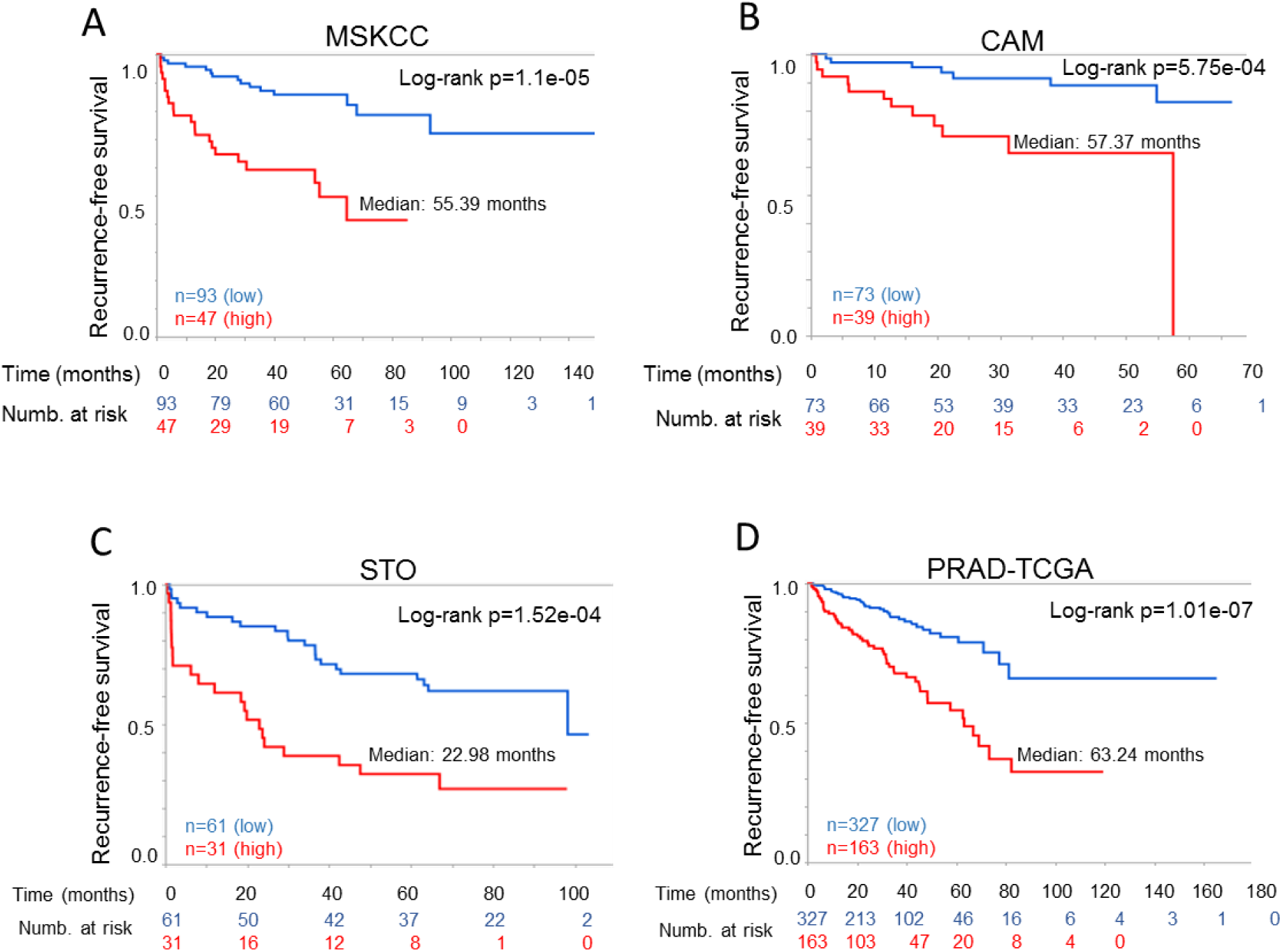
High TMCC11 expression correlates with decreased disease-free survival in several independent PCa datasets. Kaplan-Meier plots of biochemical relapse-free in the MSKCC **(A)**, Cambridge **(B)**, Stockholm **(C)** and PRAD-TGCA **(D)** datasets. Patients were divided in two categories with the upper tertile of the TMCC11 used at the cut point. Red indicates high TMCC11 group.

In Cox regression analyses, TMCC11 was a significant prognostic variable (p<0.001) with a hazard ratio (HR) of 4.1 (Table 2). In multivariate analysis, and a model constructed using a forward stepwise selection process coupled with Cox proportional hazard, TMCC11 remained a significant prognostic variable with a HR of 2.27 and 2.35 respectively (Table 2). The final model also selected pathological T-score and surgical Gleason score as significant predictors of BCR (Table 2).

**Table 2.** Uni- and multivariate Cox regression analysis of disease recurrence in several PCa datasets. Gleason – High (≥4+3): Low (≤3+4); PSA – High (≥10):Low(<10); Path Stage –High(≥T3):Low(≤T2); Positive surgical margins −Y:N; Extracapsular extension (ECE) – Y:N.

|  |  | UNIVARIATE ANALYSIS |  | MULTIVARIATE ANALYSIS |  | FINAL MODEL |  |
| --- | --- | --- | --- | --- | --- | --- | --- |
|  |  | HR (95% CI) | p-value | HR (95% CI) | p-value | HR (95% CI) | p-value |
| **MSKCC** | TMCC11 | 4.10(2.08, 8.1) | <0.001 | 2.27(1.05,4.91) | 0.038 | 2.35(1.14,4.87) | 0.021 |
|  | Biop. Gleason | 4.26(1.93, 9.42) | <0.001 | 0.96(0.36,2.59) | 0.943 |  |  |
|  | PSA | 2.98(1.53, 5.83) | 0.001 | 1.66(0.79,3.49) | 0.372 |  |  |
|  | Path Stage | 4.62(2.30,9.27) | <0.001 | 4.15(1.56,11.04) | 0.459 | 2.99(1.42,6.29) | 0.004 |
|  | Surg. margins | 2.07(1.06, 4.06) | 0.034 | 0.90(0.42,1.95) | 0.776 |  |  |
|  | ECE | 2.10(0.92,4.80) | 0.79 | 0.42(0.12,1.29) | 0.415 |  |  |
|  | Surg. Gleason | 10.56(4.87,22.86) | <0.001 | 7.92(3.16,19.87) | <0.001 | 6.78(2.92,15.73) | <0.001 |
| **CAM** | TMCC11 | 4.76(1.80, 12.59) | 0.002 | 4.28(1.53, 11.99) | 0.006 | 3.53(1.31,9.51) | 0.013 |
|  | Biop. Gleason | 3.25(1.32, 8.02) | 0.011 | 1.10(0.25,4.93) | 0.897 |  |  |
|  | PSA | 1.50(0.57, 3.95) | 0.412 | 2.82(1.00,7.98) | 0.050 |  |  |
|  | Surg. Gleason | 4.68(1.88,11.63) | <0.001 | 5.12(1.07, 24.55) | 0.041 | 4.31(1.70,10.91) | 0.002 |
|  | Surg. margins | 1.64(0.62, 4.35) | 0.324 | 2.08(0.76,5.71) | 0.156 |  |  |
|  | ECE | 1.82(0.60,5.48) | 0.288 | 0.85(0.28,2.64) | 0.780 |  |  |
| **STO** | TMCC11 | 3.00(1.65,5.44) | <0.001 | 2.69(1.46,6.11) | 0.003 | 2.89(1.56,5.36) | <0.001 |
|  | Preop. Gleason | 2.67(1.44,4.96) | 0.002 | 1.40(0.66, 2.99) | 0.381 | 2.12(1.12,4.02) | 0.021 |
|  | PSA | 1.61(0.89,2.92) | 0.116 | 1.05(0.56,1.98) | 0.879 |  |  |
|  | Surg. Gleason | 3.62(2.00,6.58) | <0.001 | 1.77(0.84,3.74) | 0.136 |  |  |
|  | Surg. margins | 1.99(1.10,3.59) | 0.023 | 1.84(0.96,3.54) | 0.068 |  |  |
|  | ECE | 4.21(2.19,8.09) | <.001 | 2.98(1.46,6.11) | 0.003 | 3.69(1.89,7.20) | <0.001 |
| **TCGA** | TMCC11 | 2.94(1.94, 4.46) | <0.0001 | 1.96(1.26,3.05) | 0.003 | 1.96(1.26,3.05) | 0.003 |
|  | Gleason | 4.08(2.27, 7.34) | <0.0001 | 2.29(1.20, 4.38) | 0.012 | 2.29(1.20, 4.38) | 0.012 |
|  | Path Stage | 3.68 (2.07, 6.51) | <0.0001 | 2.25(1.22, 4.15) | 0.010 | 2.25(1.22, 4.15) | 0.010 |

We validated the prognostic findings in additional independent publically available datasets (see Table 1, Additional File 1, Table S1 and Figure S7 for descriptions). Survival analysis demonstrated that TMCC11 was a significant (log-rank p= 5.75e-05, p= 1.52e-04 and p= P=8.03e-08) predictor of outcome in the Cambridge (CAM; n=112; (34)), Stockholm (STO; n=92; (34)) and PRAD TCGA (n=490) cohorts, segregating patients with better/worse prognosis based on recurrence data over 60, 100 and 180 months respectively (Figures 3B-3D). Results using multivariate Cox regression analysis including expression level of the TMCC11 signature and several clinical variables, demonstrate that the TMCC11 signature is an independent predictor of recurrence after prostatectomy in these datasets (Table 2). Taken together, these data suggest that the TMCC11 signature is prognostic for risk of disease recurrence after radical prostatectomy, and has an added benefit in the context of standard clinical variables in several independent datasets.

The prognostic value of the TMCC11 signature was further evident using C-statistics (Additional File 1, Table S5). The TMCC11 signature was a significant predictor across all datasets. In the TCGA-PRAD, it performed better (C-index, 0.64; confidence interval, 0.58-0.70; p<0.001) than Gleason (C-index, 0.62; confidence interval, 0.58-0.67; p<0.001) or pathological score (C-index, 0.61; confidence interval, 0.57-0.66; p<0.001). Moreover, in all the datasets, the TMCC11 signature significantly improved prognostic ability when combined with other clinical variables (Additional File 1, Table S5). The persistence of the interaction terms as significant effects proves that the TMCC11 predictive effectiveness might vary with the levels of the other clinical variables.

In selected patients from the MSKCC and TCGA-PRAD datasets with high pathological T (≥ T3) or Gleason (≥ 4+3) scores, high TMCC11 significantly stratified men at risk for disease recurrence/progression (Additional File 1, Figures S9 and S10). TMCC11 provides prognostic information in high-risk patients beyond that provided by established clinico-pathological prognostic features as demonstrated using multivariate analysis (Additional File 1, Tables in figures S9 and S10). These results suggest that TMCC11 has prognostic value in men with high-grade tumors, after RP. TMCC11 failed to stratify patients with low surgical Gleason score, however, preliminary data using the MSKCC (46) and Stockholm (34) datasets indicate that TMCC11 can stratify patients presenting with low biopsy Gleason score, suggesting that the signature may be informative for PCa management after a positive biopsy (Additional File 1, Figure S11).

### Prognostic assessment of the TMCC11 gene signature

Several gene signatures have predictive capabilities in PCa. We therefore conducted additional tests to determine the value of the TMCC11 signature when compared to other signatures, using the Bioconductor package SigCheck (55). This software allows comparison of a gene signature prognostic performance against random and known gene signatures. Initially, we analyzed the prognostic power (based on time to recurrence) of TMCC11 and other previously identified oncogenic signatures: 6 signatures for PCa (22, 25, 34, 56-58), 189 oncogenic signatures from multiple cancer types in MSigDB, and 48 breast oncogenic signatures (compiled in (59)) (n=243, Tables 3 and Additional File 1, Table S6). TMCC11 outperformed most signatures (Additional File 1, Table S6). Considering just the six PCa gene signatures (Table 3), only the Cuzick (n=31) signature achieved comparable performance to the TMCC11 across the three datasets for identifying patients with shorter time to biochemical relapse, and the performance depended on the dataset utilized. Of note, 5 genes within the Cuzick set overlap with the TMCC11 set. We obtained similar results using two other TMCC11 derived signatures, TMCC13 and TMCC3 (Table S6 Additional File 1). TMCC13 is a modified form of TMCC11 including two additional genes, E2F7 and GSG2, while TMCC3 consisted of only 3 from the TMCC11 signature that do not overlap with the Cuzick signature. These results underscore the independent prognostic value of the genes included in the TMCC11 signature.

**Table 3.**
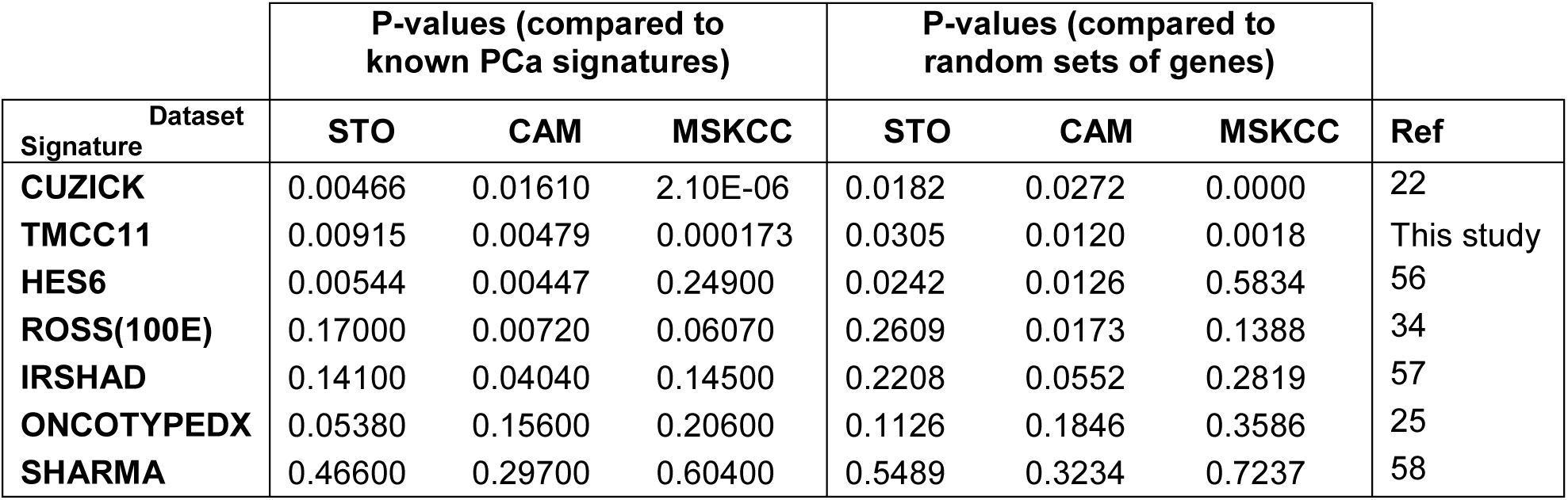
Prognostic potential of PCa signatures. Left columns: Comparative TMCC11 and known PCa signatures prognostic potential. Performance scored by log-rank test p-value of BCR difference between high and low risk groups defined by overall gene expression signature. Right columns: Comparative analysis for TMCC11 and known PCa signatures over performance against random signatures. For each signature, 10000 equal size signatures were generated at random and evaluated for predicting early relapse by log-rank test p-value. An overall bootstrap p-value score was computed as proportion of random signatures performing better than the initial signature. For both analyses, the data is sorted by first principal component of the individual rankings of the 3 columns corresponding to the Cambridge, Stockholm and MSKCC datasets. The Ross(100E) signature corresponds to the genes selected based on transcriptome profiling only. See also tables S5 and S6 in Additional File 1 for a full list with additional signatures.

| Dataset<br>Signature | P-values (compared to<br>known PCa signatures) |  |  | P-values (compared to<br>random sets of genes) |  |  | Ref |
| --- | --- | --- | --- | --- | --- | --- | --- |
|  | STO | CAM | MSKCC | STO | CAM | MSKCC |  |
| **CUZICK** | 0.00466 | 0.01610 | 2.10E-06 | 0.0182 | 0.0272 | 0.0000 | 22 |
| **TMCC11** | 0.00915 | 0.00479 | 0.000173 | 0.0305 | 0.0120 | 0.0018 | This study |
| **HES6** | 0.00544 | 0.00447 | 0.24900 | 0.0242 | 0.0126 | 0.5834 | 56 |
| **ROSS(100E)** | 0.17000 | 0.00720 | 0.06070 | 0.2609 | 0.0173 | 0.1388 | 34 |
| **IRSHAD** | 0.14100 | 0.04040 | 0.14500 | 0.2208 | 0.0552 | 0.2819 | 57 |
| **ONCOTYPEDX** | 0.05380 | 0.15600 | 0.20600 | 0.1126 | 0.1846 | 0.3586 | 25 |
| **SHARMA** | 0.46600 | 0.29700 | 0.60400 | 0.5489 | 0.3234 | 0.7237 | 58 |

We then analyzed the performance of the oncogenic signatures against 10,000 signatures consisting of the same number of genes (for the specified signature) selected at random (Tables 3 and Additional File 1, Table S7). The TMCC11 signature performed in the 97^th^ and 99^th^ percentiles, with only 3%, 1.2% and 0.18% of the random signatures demonstrating an equal or smaller p-value (empirical p-values of p=0.0305, p=0.012 and p=0.0018) in predicting relapse in the Stockholm, Cambridge and MSKCC datasets respectively. Considering the PCa signatures, only the Cuzick (n=31) signature achieved comparable performance to the TMCC11 across the three datasets (Table 3). TMCC11, TMCC13 and TMCC3 outperformed most of the oncogenic signatures described above (n=243), when tested against random signatures (Table S7 Additional File 1).

## DISCUSSION

Here, we have identified an 11-gene prognostic signature (TMCC11) for PCa progression consisting of genes associated with cell-cycle and DNA damage response. The prognostic value of this signature was confirmed on several publically available cohorts totaling 834 samples from geographically different cohorts of patients that underwent RP. TMCC11 is an independent predictor of biochemical recurrence after RP and added significant prognostic value to standard clinicopathological variables. In multivariate analysis TMCC11 was the only variable consistently predictive of disease recurrence in all of the datasets, and it significantly increased risk prediction over other clinical variables and when combined with other variables (Tables 2 and Additional File S5). Moreover, in subsets of patients with high Gleason or pathological scores, the TMCC11 signature provided a statistically significant stratification of patients identifying high and low risk groups of BCR, and preliminary data suggests that TMCC11 can stratify patients that present with low biopsy or pre-operative Gleason scores. All together, these results suggest that TMCC11 may provide relevant prognostic information in several clinical scenarios and have an impact not only on the decision of whether to provide adjuvant therapy after RP, but also on treatment management after a positive biopsy.

Genomic and transcriptomic analyses have provided insight into the complexity of prostate tumors and the existence of molecular subtypes. However, the clinical applicability of these classifications has been thwarted, due in part to the highly heterogeneous nature of PCa and the difficulty of identifying additional relevant alterations that occur at low frequencies (11-18) (60). We hypothesized that heterogeneously expressed genes can expose unidentified molecular subclasses of PCa and/or identify translationally relevant gene sets. Expression of *Tmeff2*, an androgen regulated gene, is highly variable across several different PCa datasets ((34) and Table S3, Additional File 1). Low *TMEFF2* expression significantly associated with shorter time to post-RP BCR. Although the prognostic value of low *TMEFF2* mRNA levels is uncertain, low *TMEFF2* mRNA correlates with increased androgen response of the cell cycle genes that define the TMCC11 signature in cell lines, and increased mRNA levels of the same genes in clinical datasets. Interestingly, SPINK1 also demonstrates highly variable expression across the same datasets (Additional File 1, Table S3). SPINK1 is an androgen-regulated gene highly overexpressed in approximately 10% of PCa cases (61-63). While the prognostic role of SPINK1 for PCa is unclear (64), it has been suggested that pathways downstream of SPINK1 may have translational and prognostic significance (64, 65). These observations hint to highly variably expressed genes as a potential source of information with translational value.

Currently several tissue-based genomic biomarkers offer prognostic information for patients with PCa either before or after treatment (23). The Decipher™ (24), Oncotype DX^®^ (25) and Prolaris^®^ (22) are commercially available panels based on measurement of gene expression changes at the RNA level. The Prolaris^®^ panel, based on the set described in Cuzick (22), examines the expression of 31 genes involved in cell cycle progression and 5 out of the 11 genes in TMCC11 are common to this panel. We observed a similar prognostic performance for the Cuzick (22) and the TMCC11 signatures when compared against random size-matched signatures. In addition, the prognostic power (based on p-value) of our signature vs. Cuzick (22) was dependent on the dataset utilized, but they were similarly informative and both behaved as strong risk predictors. While these comparisons need to be verified in independent studies, TMCC11 represents a smaller and more focused distinct gene set with potentially added value in specific patient subsets. The smaller size of the TMCC11 signature (11 genes vs. 31 of Cuzick (22)) is an advantage in clinical use since smaller signatures are more amenable to testing with reduced RNA quantities (i.e. biopsy samples) or even assayed with immunohistochemistry. In addition, TMCC3, a signature consisting of three genes selected from the TMCC11 signature, and that does not overlap with the Cuzick gene set, demonstrated excellent prognostic ability in SigCheck analysis. This suggests that subsets of the TMCC11 genes can be of prognostic value. Finally, the fact that our studies have independently led to the identification of a cell-cycle based signature validates the results and points to the value of using cell cycle genes as prognostic markers in PCa.

## CONCLUSIONS

Using an unconventional approach, we have identified an 11-gene signature consisting of functionally related nuclear genes with roles in DNA replication/ repair and/or cell cycle that can improve accuracy of prognosis in patients with PCa after RP in the context of current clinicopathological variables. Prognostic gene signatures containing, or based on, cell cycle gene expression changes have been identified using other approaches and different sample types. This observation not only validates our results, but also suggests that heterogeneity may lead to similar cellular consequences, providing cell cycle based signatures with rather global predictive values. The TMCC11 signature requires further validation in multi-institutional cohorts and clinical trials. In addition, the ability of TMCC11 to provide prognostic information using biopsy samples needs to be further explored.

## ABBREVIATIONS

PCa: Prostate cancer
RP: Radical prostatectomy
qRT-PCR: Quantitative reverse-transcription polymerase chain reaction
TMEFF2: Transmembrane protein with EGF like and two follistatin domains 2
AR: Androgen receptor
BCR: Biochemical recurrence
PSA: Prostate specific antigen
DFS: Disease free survival
CRPC: Castration resistant prostate cancer
CPM: counts per million
DEG: differentially expressed gene
FDR: False discovery rate

## ACKNOWLEDGEMENTS

The results published here are in part based upon data generated by The Cancer Genome Atlas Research Network (http://cancergenome.nih.gov/). RNA-Seq analysis help was provided by the Oklahoma Medical Research Foundation Quantitative Analysis Core and supported by COBRE grant 1P30GM110766-01. Histology, immunohistochemistry, slide scanning, and image analysis were provided by the Stephenson Cancer Center at the University of Oklahoma, an Institutional Development Award (IDeA) from the National Institute of General Medical Sciences (P20 GM103639) and a Cancer Center Supporting Grant Award from the National Cancer Institute (P30 CA225520), both from the National Institute of Health. Special thanks to all the patients that provided samples that made possible this study.

Supplementary information accompanies the paper.

